# Unraveling the mechanism of proton translocation in the extracellular half-channel of bacteriorhodopsin

**DOI:** 10.1101/031609

**Authors:** Xiaoxia Ge, M. R. Gunner

## Abstract

Bacteriorhodopsin, a light activated protein that creates a proton gradient in halobacteria, has long served as a simple model of proton pumps. Within bacteriorhodopsin, several key sites undergo protonation changes during the photocycle, moving protons from the higher pH cytoplasm to the lower pH extracellular side. The mechanism underlying the long-range proton translocation between the central (the retinal Schiff base SB216, D85 and D212) and exit clusters (E194 and E204) remains elusive. To obtain a dynamic view of the key factors controlling proton translocation, a systematic study using molecular dynamics simulation was performed for eight bacteriorhodopsin models varying in retinal isomer and protonation of the SB216, D85, D212 and E204. The side-chain orientation of R82 is determined primarily by the protonation states of the residues in the EC. The side-chain reorientation of R82 modulates the hydrogen-bond network and consequently possible pathways of proton transfer. Quantum mechanical intrinsic reaction coordinate calculations of proton-transfer in the methyl guanidinium-hydronium-hydroxide model system show that proton transfer via a guanidinium group requires an initial geometry permitting proton donation and acceptance by the same amine. In all the models, R82 can form proton wires with both the CC and the EC connected by the same amine. Alternatively, rare proton wires for proton transfer from the CC to the EC without involving R82 were found in an O’ state where the proton on D85 is transferred to D212.

## Introduction

As a major form of energy storage for biological organisms, the transmembrane electrochemical proton gradient is generated either through vectorial redox reactions or transmembrane proton transfer reactions.^1^ Bacteriorhodopsin, the simplest and most studied proton pump, moves protons from low (N-side) to high (P-side) concentration by harnessing light energy, creating a transmembrane electrochemical gradient.^2^–^4^ The proton-pumping reaction cycle of bacteriorhodopsin starts after light-induced all-*trans* to 13-*cis* isomerization of the K216-attached retinal Schiff Base (SB216). Following a series of proton transfer reactions, re-isomerization of retinal to all-*trans* configuration returns the protein to the ground state. The reaction cycle passes through the intermediate states bR (ground), K, L, M1, M2, N, N’ and O (Figure 1), which have been identified by time resolved spectroscopy.^5^–^7^

**Figure 1.**
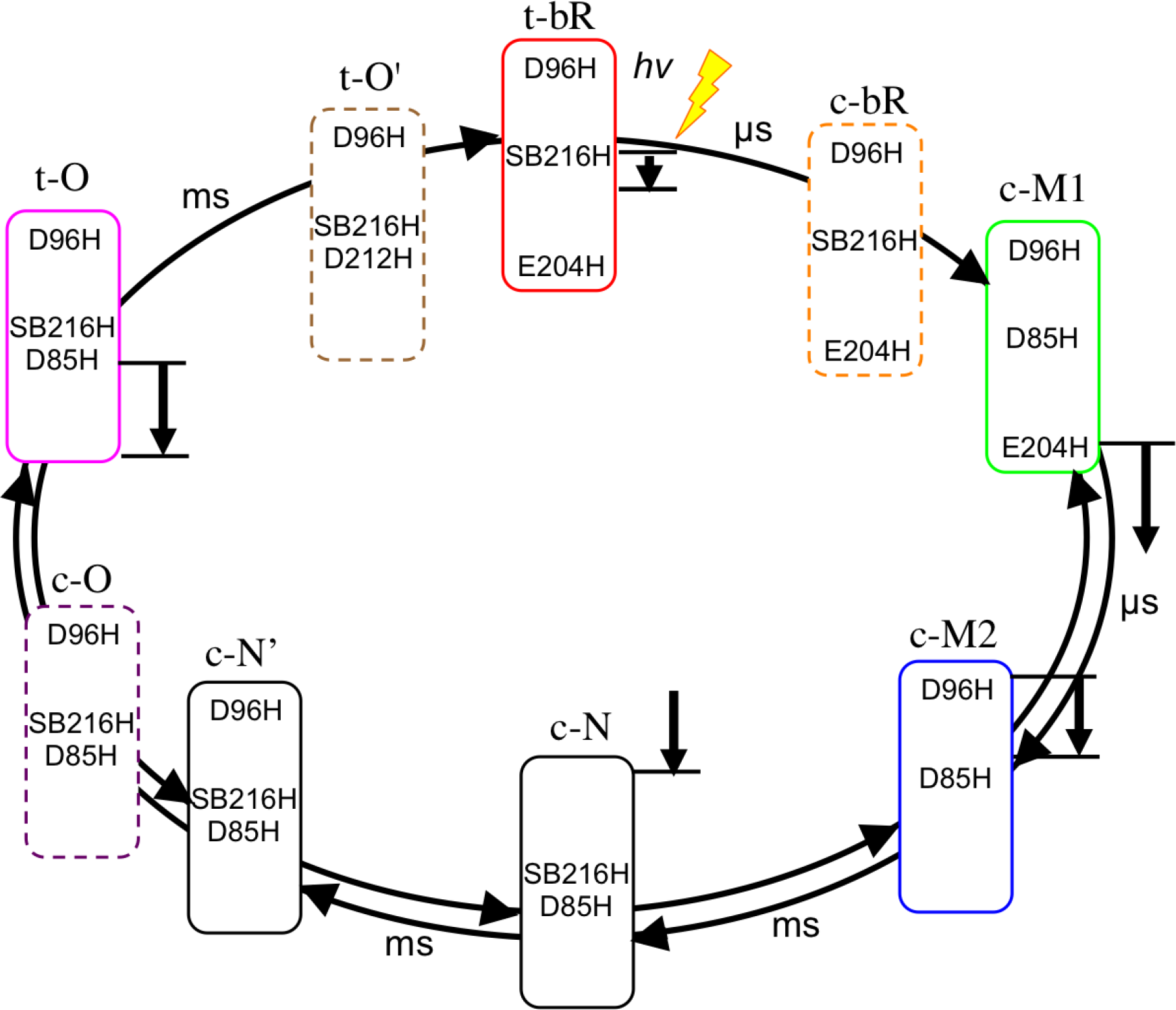
Bacteriorhodopsin photocycle. Each rectangular box represents a real intermediate states (solid border) or hypothetical states (dashed border). The box colors are used consistently in other figures to denote the individual states. Protonated residues along the proton transfer pathway are listed in the boxes. Deprotonated CC (SB216, D85 and D212) and EC (E194 and E204) residues are not shown. Vertical downward pointing arrows indicate the proton transfers that will transfer a proton from the higher pH cell interior to the lower pH extracellular side. D212 is deprotonated in all intermediates except O’ where the proton on D85 is transferred to D212.

It has been established by Fourier transform infrared (FTIR) spectroscopy and resonance Raman studies^2,8–10^ that the functional intermediates represent altered protonation states of three clusters of key residues located in separated regions (Figure 2), *i.e.*, proton uptake site D96 near the cytoplasmic N-side, the central cluster (CC) consisting of SB216, D85 and D212, and the proton exit cluster (EC) composed of E194 and E204 near the extracellular P-side of the protein. An amino acid R82 located between the CC and the EC have been shown to affect the interaction of internal waters during the bR to M transition^11,12^ as well as the rate of proton release from the EC^13^ and transfer from the CC to EC.^14,15^

**Figure 2.**
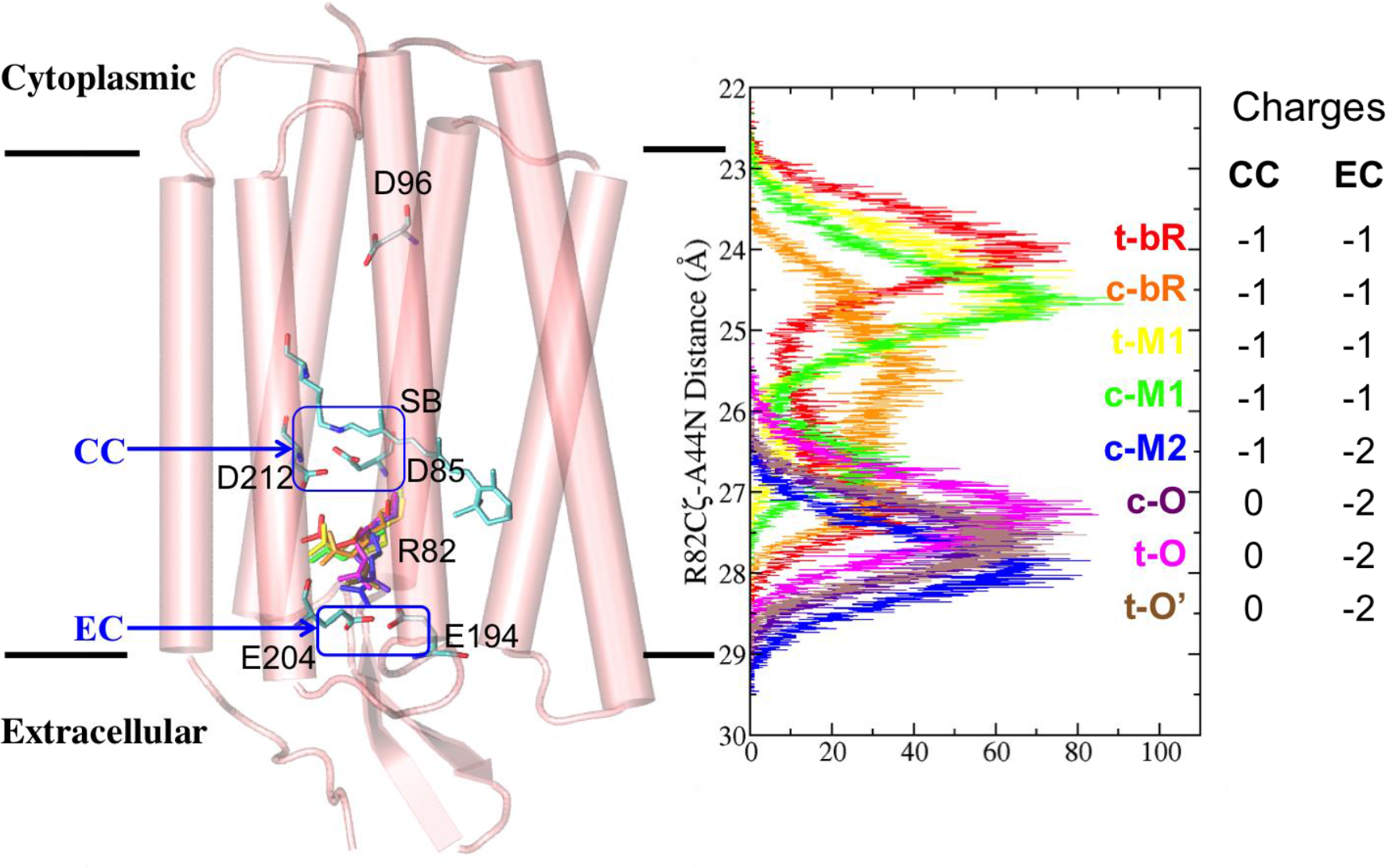
Distribution of R82 side-chain orientations in MD trajectories for different states. (Left) Monomeric structure of bacteriorhodopsin is represented by transparent cartoon in pink, protonable amino acids along the pathway are displayed by sticks in CPK color for non-hydrogen atoms based on equilibrated conformation of t-bR state model. Side-chain positions of R82 in equilibrated conformations of all seven models are shown by colored sticks: red for t-bR, orange for c-bR, yellow for t-M1, green for c-M1, blue for c-M2, purple for c-O, magenta for t-O and brown for t-O’ state. (Right) The distribution of R82Cζ-A44N distance is plotted for each simulation model. The prefix t- and c- refer to all-trans and 13-cis isomerization of retinal respectively. The color scheme and labeling of the state apply to rest of the figures. The sums of side-chains charges for the CC and the EC residues are given next to the label for each state.

A series of X-ray crystal structures^16–20^ have been solved for intermediates trapped by mutation or freezing, which provide a structural basis for studying the proton-pumping mechanism of bacteriorhodopsin. The structures show small conformational changes associated with the key residues in the CC and the EC.^2^ However a crystal structure represents a single snapshot of a dynamic biomolecule, which is influenced by its crystallization condition and other factors.^21^ Moreover, X-ray diffraction usually cannot resolve the positions of hydrogen atoms. While it is possible to monitor the kinetics of light-activated changes in bacteriorhodopsin, only a subset of the motions in the protein can be assigned to the observed optical or infrared spectral changes. Molecular dynamics (MD) simulations provide a means for viewing in atomic detail the motions of protein and water molecules to reveal key interactions that may be associated with proton pumping.

Extensive experimental and theoretical studies have been carried out on bacteriorhodopsin. The mechanism of proton uptake and transfer in the cytoplasmic half-channel (from D96 to SB216) has been studied with FTIR and molecular mechanics (MM) methods.^9,22,23^ Computational studies have been carried out to rationalize the short-range proton transfer, from SB216 to D85,^24–29^ and within the EC and nearby R82^7,30,31^ using quantum mechanical (QM) calculation or QM/MM hybrid methods.

The longest proton-transfer path in bacteriorhodopsin is between the CC and the EC, a region that spans ∽13 Å as measured by the distance between the carboxylate oxygens of D85 and E204. This long-range proton translocation takes place during the O to bR transition, the final step of the proton-pumping cycle. Despite decades of studies, molecular details of how a proton moves between the CC and the EC are still not fully understood. Proton translocation in this region is also associated with protein conformational change and water relocation. Different models have emerged^14,15,32,33^ based on two main hypotheses. In *accessibility-switch* models, proton transfer pathways in different regions of the protein are opened and closed during the reaction cycle to facilitate the forward transfer and block the back transfer. Alternatively, *affinity-switch* models focus on changes in proton affinity of key residues along the transport channel during the photocycle. Hence, a “wrong-way” proton transfer would not occur because of the proton affinity changes of the key residues. In accessibility-switch models, Lanyi *et al*. have proposed that protons are transferred through a hydrogen bonded network connected by R82;^14^ Kandt *et al*. suggest that the movement of R82 allows a direct and continuous Grotthus-type proton transfer pathway from SB216 to the EC.^15^ By contrast, time-resolved FT-IR spectroscopy and kinetic analysis by Lorenz-Fonfria *et al*.^32^ and continuum electrostatics calculations of the energetics of proton transfer by Onufriev *et al*.^33^ support a mechanism of proton transport via a switch that is based on changes in the proton affinity of key residues rather than accessibility.

This paper reports a computational study of the bacteriorhodopsin monomer in different functional states. To examine the relative importance of retinal isomerization and residue protonation states in determining the R82 position and proton wire connectivity, atomic simulation models containing 13-*cis* (c) or all-*trans* (t) retinal with the same protonation states were studied. Eight computational models including four experimentally characterized intermediate states t-bR, c-M1, c-M2, and t-O as well as four hypothetical states bR-like (c-bR), M1-like (t-M1), O-like (cO) and O’ (t-O’) were constructed with explicit water and membrane (Table 1 and Figure 1). MD simulations in conjunction with hydrogen bond (H-bond) analysis and QM intrinsic reaction coordinate (IRC) calculations were applied to investigate the proton transfer mechanism in the extracellular half-channel of bacteriorhodopsin. A correlation was identified between the side-chain position of R82 and the protonation states of E194 and E204 in the EC. The R82 side-chain orients toward the EC when E194 and E204 are both deprotonated (M2 and O states), while it oscillates between the CC and the EC when E194 forms a H-bond with a protonated E204 (bR and M1 states). The R82 position shifts slightly downward when the retinal is 13-*cis* compared with its all-*trans* counterpart. The IRC paths indicate that proton transfer through the R82 side-chain requires an initial geometry that allows proton donation and acceptance to take place sequentially by the same amine. Proton wires from R82 to the CC and from R82 to the EC were found in all the models. A preferred CC to EC directionality of proton-transfer in M2 and O states was inferred from the higher probability of proton wires connecting R82 to the EC. Alternative pathways were also proposed for proton translocation between the CC and the EC through a Grotthuss-type transfer without involving R82. The Grotthuss pathways are rare and only occur in one direction from the CC to the EC in c-M2, c-O state, and more likely in a transient t-O’ state, where the proton has transferred from D85 to E212 late in the reaction cycle.

**Table 1.** Simulation models constructed in this study.

| Models | Starting structures | Schiff base configuration | R82 orientation | Protonated residues | Charges |  |
| --- | --- | --- | --- | --- | --- | --- |
|  |  |  |  |  | CC | EC |
| **t-bR** | <u>1QHJ</u> | 13-trans | CC | D96, D115, SB, E204 | -1 | -1 |
| c-bR | 1P8H | 13-cis | CC | D96, D115, SB, E204 | -1 | -1 |
| **c-M1** | <u>1P8H</u> | 13-cis | CC | D96, D115, D85, E204 | -1 | -1 |
| t-M1 | 1QHJ | 13-trans | CC | D96, D115, D85, E204 | -1 | -1 |
| **c-M2** | <u>1CWQ</u> | 13-cis | EC | D96, D115, D85 | -1 | -2 |
| c-O | 1P8H | 13-cis | CC | D96, D115, SB, D85 | 0 | -2 |
| **t-O** | 1QHJ | 13-trans | CC | D96, D115, SB, D85 | 0 | -2 |
| t-O' | 1QHJ | 13-trans | CC | D96, D115, SB, D212 | 0 | -2 |

## Materials and Methods

**MD simulation models of different bacteriorhodopsin states.** To expedite equilibration, crystal-structures of the ground state with an all-*trans* retinal and early M1 state with a 13-*cis* retinal were adopted as starting structures. Considering the distortion observed on helix F backbone in the M2 state relative to other states, a structure trapped in the M2 state was also selected. High-resolution crystal structures of wild-type bacteriorhodopsin trapped in three intermediate states bR (PDB ID: 1QHJ^16^), M1 (1P8H^17^ model 1) and M2 (1CWQ^18^ chain B) were used to build the simulation models (Table 1) corresponding to four intermediate states bR, M1, M2, and O as well as four hypothetical states c-bR (with 13-*cis* retinal), t-M1 (with all-*trans* retinal), c-O (with 13-*cis* retinal) and t-O’ (D212 is protonated by D85). As no O state structure is available at neutral pH, a t-O model was built based on the t-bR structure. All simulation models were constructed as monomers including residue number 1 to 231 since the monomeric bacteriorhodopsin has been demonstrated to function with a proton-pumping efficiency comparable to the native trimer.^34^ The missing N-terminal residues number 1 to 4 were built with PyMol,^35^ the missing loop residues number 157 to 161 were constructed using the bR state crystal structure 1QHJ as a template. The constructed atoms were relaxed in vacuum with the rest of the protein atoms fixed.

Unbiased all-atom MD simulations were carried out with NAMD version 2.9.^36^ The CHARMM36 all-atom force field^37^ was used for protein, lipids ions and retinal (see the Supporting Information). Water molecules were described explicitly using the TIP3P model.^38^ Crystallographic water molecules were retained in the simulations. Protonation states of the key residues were assigned for each model based on the literature^5,6,39^ as described in Table 1. The protein was first embedded in a bilayer of 164 POPC, solvated and neutralized by Na^+^ and Cl^-^ ions to reach a salt concentration of 1 M, mimicking the biological conditions of halobacteria. Depending on the initial structure, each model contains about 16400 water molecules. Periodic boundary conditions were applied with a box size of 76 × 90 × 120 Å^3^. The system first underwent Langevin dynamics at constant volume. While keeping rest of the system fixed, the lipid tails were melted by performing 1000 steps of conjugate gradient energy minimization followed by 0.5 ns simulation. The system was then subjected to 1000 steps of minimization and 0.5 ns equilibration with the protein restrained with a 1 kcal/mol/Å^2^ harmonic force constant. After removal of harmonic constraints, the system was equilibrated for another 0.5 ns. The production simulations were performed using constant area-isothermal-isobaric algorithm with a temperature of 298 K and a pressure of 1.0132 bar. A time step of 2 fs was used throughout the simulations. A cutoff of 12 Å and a switching distance of 10 Å were applied for non-bonded interactions. Electrostatics was evaluated with the particle mesh Ewald method. The SHAKE algorithm^40^ was used to constrain all bonds involving hydrogen atoms. To enhance the conformational sampling, we performed four independent 20 ns simulations after 1.5 ns of constrained simulation and relaxation reaching a total simulation time of 81.5 ns for each bacteriorhodopsin model. However, given the modest time scale, these simulations serve to sample the proton transfer networks that would be found in each protonation intermediate rather than explore the conformational dynamics between intermediates. The MD trajectories reached equilibrium within the first 10 ns. The analysis was performed based on the last 10 ns of the 20 ns simulation for each model.

**Water occupancy between the CC and the EC.** Protein-water interactions modulate the pattern of the intra-protein H-bond network and potentially play a role in the proton transfer between the CC and the EC. VolMap, a VMD^41^ plugin that calculates the average density and creates volumetric map based on the atomic coordinates and properties of selected atoms, has been widely used to study many biological systems.^42–44^ To determine which water positions between the CC and the EC are most stable over the course of simulations, the trajectories were first centered and RMS fitted to the equilibrium conformation from the last frame of the trajectory. The average occupancy of the water oxygens between the CC and the EC were generated with the VolMap plugin. The waters at higher-occupancy positions during simulation were considered capable of forming more stable H-bonds with nearby residues.

**H-bond analysis for protein side-chains.** The H-bonds between side-chains of key residues in the extracellular half-channel were identified using the HBond plugin of VMD.^41^ Based on the van de Waal radii reported by Bondi,^45^ Alvarez,^46^ as well as H-bonds categorized by Jeffery,^47^ the H-bond criteria include: 1) the distance between the hydrogen donor and acceptor atoms is within 3.5 Å; 2) the angle formed by donor-hydrogen-acceptor is less than 140°.^41^ The probability of each H-bond pair was then calculated.

**Proton wires connecting the CC and the EC**. The Grotthuss mechanism^48,49^ describes how an “excess” proton or protonic defect can diffuse through a H-bond network of water molecules or other H-bonded liquids through formation and cleavage of covalent bonds. In biological systems, the Grotthuss mechanism suggests that proton transfer can occur when proton wires are formed by water and polar side-chains of protein residues. For each frame of the MD trajectories, a list of H-bond pairs of polar side-chains and water molecules in the CC-EC region were first generated. H-bond paths between a pair of predefined donor and acceptor atoms that are far apart were constructed by threading the intervening H-bonded pairs in each simulation frame. The equilibrium probability of forming a particular proton wire in 20,000 total simulation frames was calculated for each model.

**Quantum calculations of proton transfer pathway near an arginine side-chain.** The Grotthuss mechanism relies on the sites that can both donate and accept protons. The protonated Arg has no lone pairs and must donate a proton before it can accept one. To study the possible role of an Arg side chain in facilitating proton transfer, quantum mechanical IRC calculations were performed on a simplified model containing only a methyl guanidinium, hydronium and hydroxide ions in the gas phase. A hydroxide ion was first placed in different positions around each amine N to mimic the H-bond geometry between water and R82. Then a hydronium ion was assigned a random position in the vicinity of each amine N but beyond the distance of covalent bonding. The minimum energy paths for proton transfer from the hydronium to the hydroxide with different starting geometries were calculated through the QM intrinsic reaction coordinate (IRC)^50,51^ procedure in the forward direction with Guassian09^52^ at HF/6-31G(d,p) level. The IRC method was developed to find the path for molecules to move down the product and reactant valleys with zero kinetic energy. Such a minimum energy reaction pathway is defined by mass-weighted Cartesian coordinates between the transition state of a reaction and its reactants and products. This approach has been successfully applied to study reaction pathways in many systems, such as gas-phase reaction,^53^ isomerization of serine-water clusters,^54^ and proton transfer in formamide-thioformamide dimer.^55^

## Results and Discussion

This study is to investigate the potential pathways for proton translocation in the extracellular half-channel of bacteriorhodopsin. Protons are transferred from the higher pH cytoplasm to the lower pH extracellular side (Figure 1 and 2). Previously pKa calculations, starting with the crystal structures of bacteriorhodopsin trapped in different intermediate states, have been shown to favor the metastable protonation states expected for each intermediate.^33,39,56^-^58^ The current work explores the changes in accessibility of proton transfer in various protonation and retinal isomeric states. In the light-activated bacteriorhodopsin, the reaction cycle is initiated by the torsional isomerization of retinal C13=C14 bond and SB216 C15=NÇ bond followed by multiple small protein motions including backbone tilting of helices C, E and F, R82 side-chain reorientation (Figure S1). Therefore, MD simulation models were created with a protein conformation close to the corresponding intermediate state to explore the H-bond connectivity in these different states rather than the transition between different intermediates. The structure of bacteriorhodopsin has reached equilibrium in these distinct intermediates after 8 ns relaxation in most of the trajectories as shown by the RMSD of the protein backbone non-hydrogen atoms as a function of time (Figure S7). Nevertheless, there is the possibility that such kind of brute force simulations did not converge within ∽20 ns. Enhanced sampling methods could alsobe applied achieve sufficient sampling and ensure convergence.

To investigate the role of retinal isomerization in pathway connectivity, hypothetical states (bR-like, M1-like and O-like) were generated by varying the isomerization state of the retinal C13=C14 bond for each protonation state. The bR-like (c-bR) model could be considered as a model of bR555, which was suggested in 13-cis, 15-syn isomerization by Bajaj *et al^59^.* The M1-like (t-M1) has the same protonation as c-M1 except for a 13*-trans, 15-anti* configuration. The O-like (c-O) model mimics a pre-O state, except for a 13-*cis*, *15-syn* configuration.

Several different force fields have been developed for the ground state retinal^60^-^62^ and reviewed in a sophisticated study by Bondar *et al^63^* Since the current study involves retinal SB with different protonation states, we chose the parameter set released in the CHARMM36 all-atom force field, which provides parameters for both protonation states of the retinal SB and is consistent with force field used for the protein and lipids. This parameter set (not published) was developed in the same manner as CHARMM General Force Field for drug-like molecules.^64^ The cofactor structure is preserved reasonably well based on the simulations of both isolated retinal and retinal within baceriorhodopsin. In different bacteriorhodopsin models, the RMSDs of retinal SB non-hydrogen atoms range from 0.3 to 0.8 Å after equilibration. The torsion angles for C14-C15=Nζ-Cε and C12-C13=C14-C15 were also preserved. As will be seen below that the H-bond connectivity outside the CC region depends more on the protonation state of key residues than the retinal isomerization. The H-bond analysis for the key residues (Table 2) and water near SB-D85-D212 indicates that SB-water402 H-bond preserved in 10∽50% of the simulation frames for c-BR, c-M1, t-M1, c-O and t-O models, but not significant for t-BR, c-M1 and t-O’ models. The active site waters 400, 401 and 402 form dynamic H-bond with SB, D85 and D212.

**Table 2.** Percentage of direct side-chain to side-chain H-bonds on the proton transfer pathway between the CC and the EC found in individual MD frames.

| Donor | Acceptor | t-bR | c-bR | t-M1 | c-M1 | c-M2 | c-O | t-O | t-O' |
| --- | --- | --- | --- | --- | --- | --- | --- | --- | --- |
| **SB216** | **D85** | 100 | 4 | N/A | N/A | N/A |  |  | 92 |
| **SB216** | **D212** |  | 4 | N/A | N/A | N/A | 53 | 76 |  |
| **D85** | **SB216** | N/A | N/A | 38 | 14 |  | N/A | N/A | N/A |
| **D85** | **D212** | N/A | N/A |  |  |  |  | 80 | N/A |
| **D212** | **D85** | N/A | N/A | N/A | N/A | N/A | N/A | N/A | 37 |
| **D212** | Y57 | N/A | N/A | N/A | N/A | N/A | N/A | N/A | 7 |
| T89 | **SB216** | N/A | N/A | 3 |  | 97 | N/A | N/A | N/A |
| T89 | **D85** | 28 | 72 | 2 |  |  |  | 1 | 3 |
| Y57 | **D212** | 99 | 99 | 99 | 99 | 83 | 100 | 51 | 47 |
| W86 | **D212** |  | 38 | 9 | 24 | 96 | 92 | 55 | 6 |
| Y185 | **D212** | 75 | 97 | 99 | 99 | 99 | 84 | 96 | 86 |
| R82 | **D212** | 45 |  | 26 | 3 |  |  |  |  |
| R82 | Y57 | 23 | 18 | 29 | 39 |  |  |  |  |
| R82 | Y83 |  |  |  |  | 3 |  | 1 |  |
| R82 | ***E194*** | 22 | 19 |  |  | 94 | 82 | 39 | 46 |
| R82 | ***E204*** | 1 | 2 |  |  | 75 | 93 | 52 | 98 |
| R82 | T205 | 2 | 30 | 2 | 9 |  | 2 |  |  |
| Y83 | ***E194*** | 66 | 25 | 19 | 54 | 99 | 100 | 99 | 100 |
| ***E204*** | ***E194*** | 83 | 92 | 82 | 45 | N/A | N/A | N/A | N/A |
| ***E204*** | S193 | 1 | 1 | 4 | 7 | N/A | N/A | N/A | N/A |
| S193 | ***E204*** | 10 | 16 | 5 | 6 | 99 | 93 | 90 | 57 |
| S193 | ***E194*** |  | 15 | 38 | 16 |  |  |  |  |

H-bonds formed between the side-chains of the key residues without intervening waters. The residues in the CC are highlighted in bold and those in the EC are in bold italic style. The percentage describes the number of MD frames with a designated H-bond out of the 20,000 total frames for each model. N/A denotes that H-bond is not possible as the putative proton donor (Glu or Asp) is deprotonated or the putative proton acceptor (SB216) is in a protonated state without a lone pair of electron. An empty box indicates a bond is formed <1% of the time.

**R82 orientation is driven by electrostatic interactions with the CC and the EC.** The role of R82 in the early stage of the proton-pumping cycle has been explored previously,^65^ showing that the downward reorientation of R82 is coupled to protonation of D85. Here we use the atomic distance between R82Cζ and A44N as a metric to define the side-chain position of R82 (Figure 2 and S2). In crystal structures trapped in t-bR (PDB code 1QHJ), c-M1 (1P8H) and c-N’ (1P8U) states,^66^ the side-chain of R82 adopts an upward orientations toward the CC with a R82Cζ-A44N distance of 24.5 to 24.9 Å; while in crystal structures trapped in c-M2 (1CWQ), acid blue t-O (1X0I)^19^ and L93A mutant t-O (3VI0)^20^ intermediate states, a downward orientation toward the EC was observed with a R82Cζ-A44N distance between 26.8 and 27.5 Å. In the current work, the R82 side-chain is defined as pointing upward if the R82Cζ-A44N distance is shorter than 26 Å and downward if this distance is longer than 26 Å.

The R82 side-chain positions are considered equilibrated after first 10 ns equilibration, as can be seen by the time series of R82Cζ-A44N distance in Figure S5. The distribution of R82 positions was extracted from the MD trajectories for each protonation and retinal isomer model (Figure 2). A correlation was found between the R82 position and the charge on the CC and the EC. The states with a **-**1 charge on both the CC and the EC generally show a bimodal distribution of the R82 side-chain orientation, with the major population pointing up toward the CC and a minor population pointing down toward the EC. The states with a **-**2 charge on the EC and 0 or **-**1 charge on the CC always find R82 pointing downward. The hypothetical c-bR model has a wide distribution with upward and downward orientations and a unique, high probability to be in the middle of the CC to the EC pathway. Despite c-O, t-O and t-O’ models being built based on c-M1 and t-bR crystal structures where R82 points up toward the CC, the R82 side-chain rapidly moves downward in the unconstrained MD simulation with E194 and E204 side-chains both being deprotonated in c-M2, c-O, t-O and t-O’ states. Thus, the negative charge on the EC attracts the positively charged R82 side-chain and stabilizes its downward conformation. It should be pointed out that the electrostatic attraction between R82 and the two clusters could be potentially overstated with a non-polarizable force field, which may overestimate the population of the downward conformation when the EC charge equals -2.^67^

Besides the charge state, the retinal configuration subtly affects the R82 side-chain position. With the same combination of protonation states, 13*-cis* isomerization of retinal is associated with a slight downward shift of the R82 position compared with their *all-trans* counterparts, as can be seen by comparing t-bR with c-bR, t-M1 with c-M1 and t-O with c-O (Figure 2).

**Correlation of key residue motions and H-bond patterns.** Concerted motions play a critical role in long-range interactions in proteins as they reflect alterations in H-bond pattern caused by protonation changes of the key residues (Figure 1 and S2). The equilibrated MD trajectories show that the side-chain orientation of R82 is correlated with the H-bonds formed between residue pairs in the CC and the EC (Figure S3). Equilibrium probabilities of H-bonds between the polar side-chains in the CC-EC region are given in Table 2 for each model. Using vectorial H-bonds as edges and side-chain positions as nodes, a molecular graph was constructed for each model, depicting a H-bonded network solely composed of polar residue side-chains (Figure 3A). These H-bonds were evaluated in a pairwise fashion; thus, multiple groups do not necessarily connect in the same MD snapshot.

**Figure 3.**
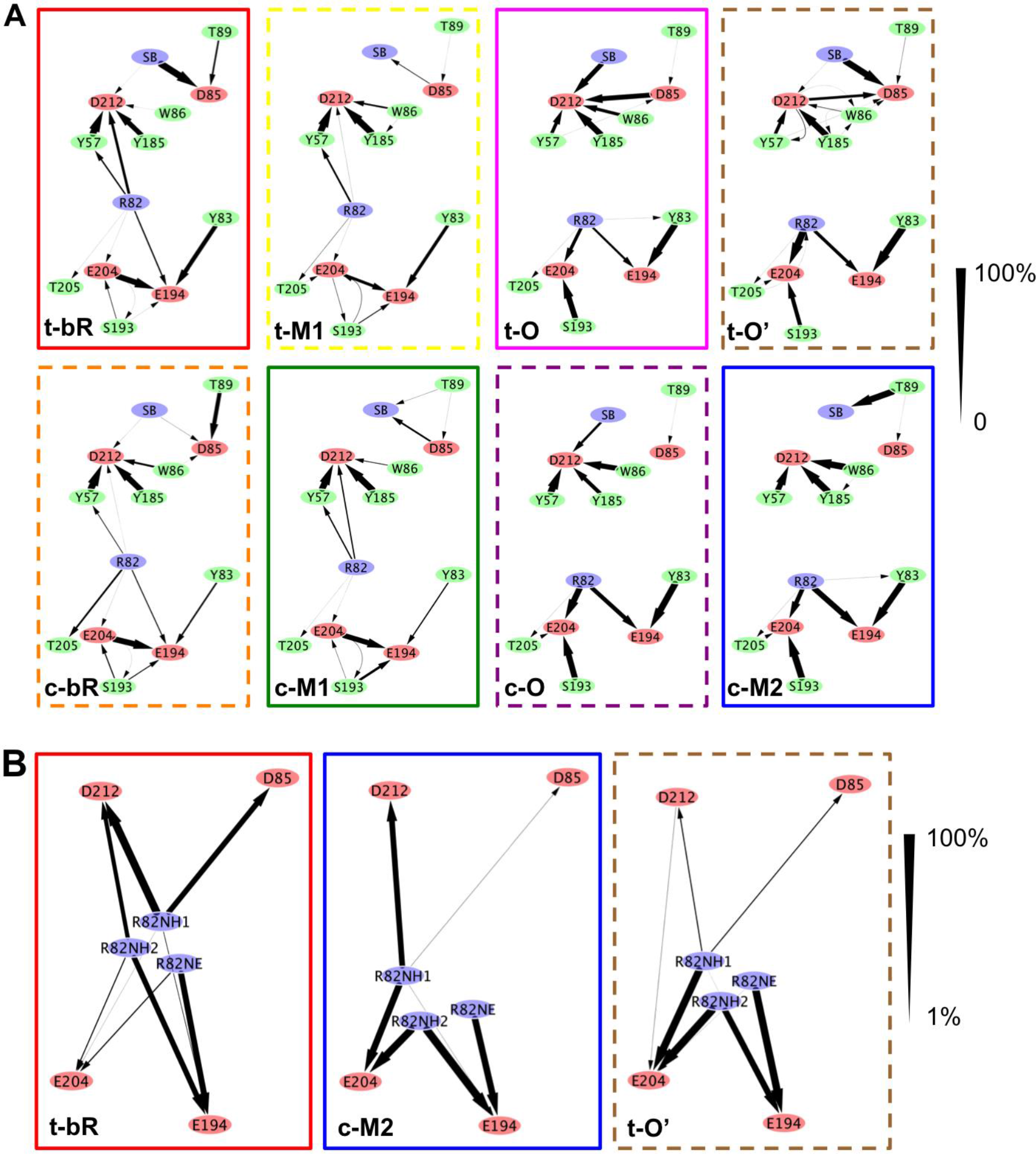
H-bond network between the CC and the EC. (A) Probabilities of H-bonds formed between protein side-chains based on the data in Table 2. Each node represents the position of a side-chain polar group defined by the X-Y coordinates of t-bR x-ray structure. Each edge represents the direction (arrow) and probability (thickness) of a single, direct H-bond. (B) Probabilities of proton wires formed in t-bR, c-M2 and t-O’ based on the data in Table 3. Only the paths with >1% probability are shown. Each edge represents a proton wire formed by waters and side-chain polar groups from the R82 side-chain to the CC or the EC with less than six sequential H-bonds, or from the CC to the EC with less than eight sequential H-bonds. The pattern found in t-bR model is similar to that of c-bR, t-M1 and c-M1 models, while the pattern found in c-M2 model is similar to that of c-O and t-O.

In the CC, as D212 and D85 are both anionic and uncoupled in the t-bR model, D212 accepts a H-bond from the R82 side-chain in 45% of the total frames, while D85 always accepts a H-bond from the Schiff Base SB216 (Table 2 and Figure S4C). Comparing the probabilities of forming a H-bond from SB216 to D85 in t-bR and c-bR or a H-bond from D85 to SB216 in t-M1, c-M1 and c-M2, it can be seen that the *all-trans* retinal facilitates the SB216-D85 connection (Table 2). Upon photoisomerization of retinal to the 13*-cis* configuration, conformational changes lead to dissociation of SB216 and D85. Due to the high pKa of 13*-cis* SB216,^39^ the proton is expected to be transferred from SB216 to D85, forming the early M state (c-M1).

In the EC, residue motions of E194 and E204 are highly correlated with the side-chain orientation of R82 (Figure S3). In t-bR, c-bR, t-M1 and c-M1 models, the protonated E204 forms a H-bond to E194 with a high probability (Table 2), regardless of whether the R82 side-chain points toward the CC or the EC. By contrast, in c-M2, c-O, t-O and t-O’ models, the deprotonated E204 and E194 move apart from each other (Figure S4E) and either can accept a H-bond from the downward orientated R82. H-bonds from Y83 to E194 and from S193 to E204 are also stabilized in the later states (Table 2).

The key event that returns the system to the ground state during the t-O to t-bR transition is proton transfer from D85 to the EC (Figure 1). The direct H-bonds among side-chains never form a complete unidirectional path connecting D85 to the EC in the simulation. However, D212 is likely to be directly connected to the R82 side-chain that can then connect to the EC (Table 2, Figure 3A). This suggests that D212 may be more accessible than D85 for proton transfer to the EC. Comparing the hypothetical c-O state with the t-O intermediate indicates that isomerization of retinal from 13*-cis* to *all-trans* allows SB216Nζ to approach D212 and D85 (Figure S4B and C). Even though the charge state of the CC and R82 orientation were the same in c-M2, c-O, t-O and t-O’, a direct H-bond between D85 and D212 was only observed when retinal is in the all-*trans* configuration (Table 2 and Figure 3A). The H-bond formed from the protonated D85 to D212 in t-O (80%) was more stable than from the protonated D212 to D85 in t-O’ (37%), The H-bond formed from SB216 to D85 was more stable in t-O’ (92%) than from SB216 to D212 in t-O (76%). Therefore, moving the proton from D85 to D212 to form t-O’ increases the hydrogen bond connectivity and thus can facilitate proton translocation from the CC to the EC.

**Table 3.** Probabilities of proton wires formed between the CC and EC residues.

| First donor | Last acceptor | t-bR | c-bR | c-M1 | t-M1 | c-M2 | c-O | t-O | t-O' |
| --- | --- | --- | --- | --- | --- | --- | --- | --- | --- |
| *E204* | *D85* | *0.0* | *0.0* | *0.0* | *0.0* | *N/A* | *N/A* | *N/A* | *N/A* |
|  | *D212* | *0.0* | *0.0* | *0.0* | *0.0* | *N/A* | *N/A* | *N/A* | *N/A* |
| *D85/D212\** | *D212/D85* | *N/A* | *N/A* | *51.0* | *33.7* | *83.3* | *92.2* | *79.8* | *83.5* |
|  | *E204* | *N/A* | *N/A* | *0.0* | *0.0* | **0.02** | **0.3** | *0.0* | **2.7** |
|  | *E194* | *N/A* | *N/A* | *0.0* | *0.0* | *0.0* | *0.0* | *0.0* | *0.0* |
| R82Nη1 | D85 | 66.3 | 85.4 | 33.9 | 20.4 | 2.9 | 0.0 | 2.6 | 11.6 |
|  | D212 | 94.3 | 96.6 | 95.4 | 94.8 | **69.8** | **64.0** | **41.7** | **12.3** |
|  | E204 | 1.5 | 0.0 | 0.0 | 0.0 | **81.4** | **89.6** | **83.1** | **91.6** |
|  | E194 | 6.3 | 0.0 | 0.0 | 0.0 | 1.9 | 0.9 | 18.6 | 1.0 |
| R82Nη2 | D85 | 0.0 | 0.0 | 0.0 | 0.0 | 0.0 | 0.0 | 0.0 | 0.0 |
|  | D212 | **50.1** | **31.5** | **47.3** | **67.5** | 0.0 | 0.0 | 0.0 | 0.0 |
|  | E204 | **11.8** | **15.0** | **19.6** | **13.9** | 90.6 | 96.0 | 84.4 | 95.7 |
|  | E194 | **65.7** | **64.4** | **49.3** | **38.5** | 97.8 | 91.9 | 91.6 | 75.8 |
| R82Nε | D85 | 0.0 | 0.0 | 0.0 | 0.0 | 0.0 | 0.0 | 0.0 | 0.0 |
|  | D212 | 0.0 | 0.0 | 0.0 | 0.0 | 0.0 | 0.0 | 0.0 | 0.0 |
|  | E204 | 13.2 | 21.3 | 23.0 | 21.0 | 0.1 | 0.2 | 25.1 | 1.3 |
|  | E194 | 72.2 | 83.6 | 74.7 | 70.2 | 81.3 | 94.6 | 71.1 | 91.3 |

Proton wires connecting key residues via waters and other polar side-chains. The percentage describes the number of MD frames with a designated proton wire out of the 20,000 total frames for each model. N/A means H-bond is not possible as the putative proton donor (Glu or Asp) is deprotonated. Connections in the top rows (noted in italics) are for proton wires that do not pass through R82. For a given side-chain N atom of R82, the percentages in bold indicate possible connections between the CC and EC residues via R82-mediated proton hopping. This is through Nη2 in the early stage and through Nη 1 in the late stage during the photocycle.
\* In t-O' state, proton wires begin from D212 instead of D85.

**Variation in the water occupancy between the CC and the EC.** Waters greatly enhance the flexibility of pathways for proton transfer by bridging the separated proton donors and acceptors.^3^ However, the energy barrier is higher in a water-mediated proton transfer than a direct transfer between the H-bonded partners.^68^ To further explore the proton transfer pathway between the CC and the EC, the dynamics and distribution of waters in this region were investigated.

Crystal structures trapped in t-bR, c-M1 and c-M2 states contain eight to nine water molecules in the CC-EC region (Figure 4). In t-bR state, these nine waters are labeled as 400, 401, 402, 403, 406, 407, 408, 409 and 427.^16^ In c-M1 state, the waters change their positions except for water 401 and 402. The structure of c-M2 has a different numbering scheme of the waters. By comparison, six positions of water 400, 401, 403, 406, 408 and 427 are relatively well conserved within the three crystal structures. The major differences are near the EC. Water positions 400, 401, 402, 406, 407 and 408 are shifted in the c-M2 structure, while water 409 was missing in c-M1 structure.

**Figure 4.**
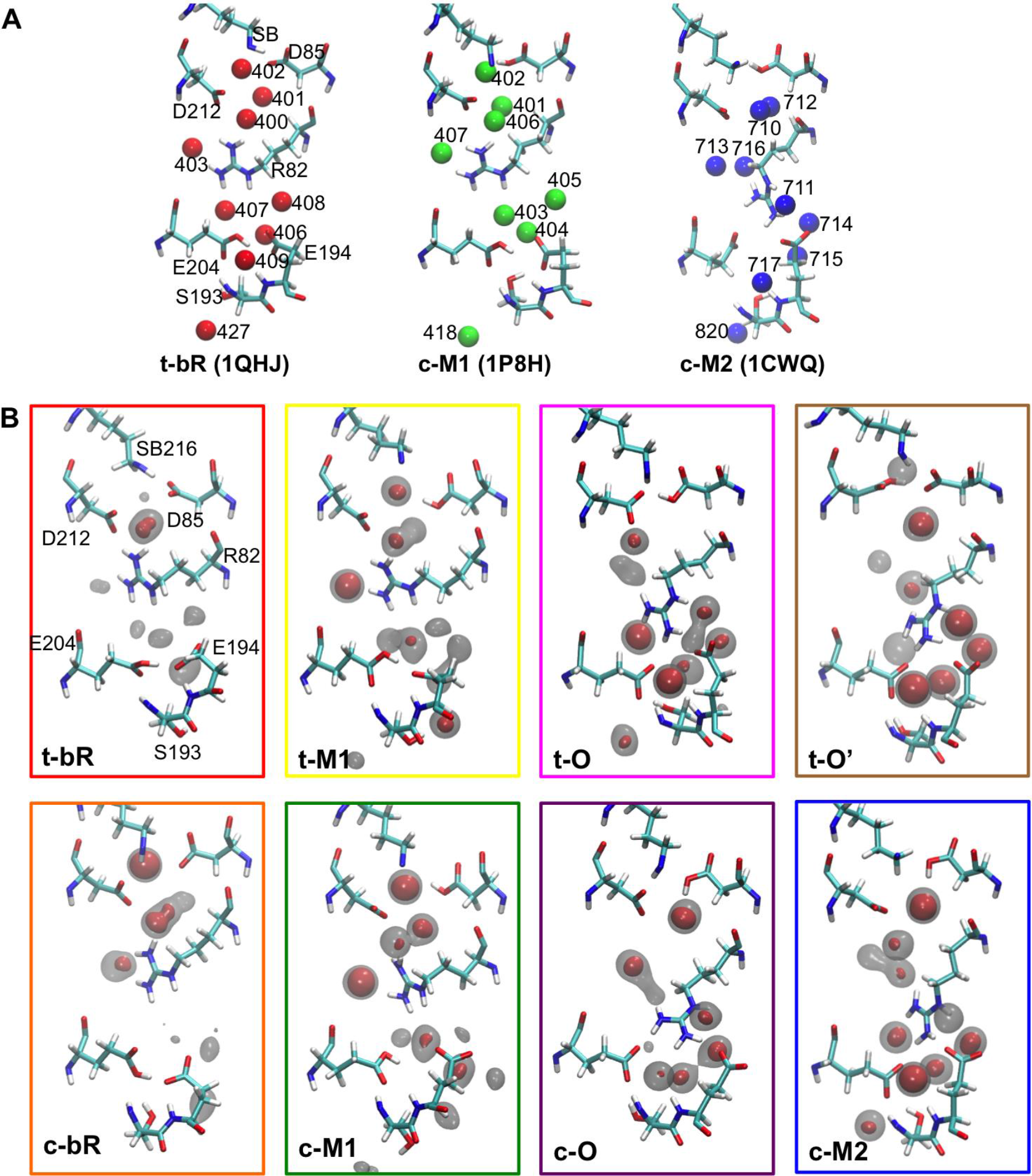
Water occupancy in bacteriorhodopsin models. (A) Crystal structures of three intermediate states: t-bR (1QHJ), c-M1 (1P8H) and c-M2 (1CWQ) with water oxygen positions displayed by color spheres, protein by licorice, all residues and water oxygens are labeled based on the PDB files. (B) Occupancy maps for water oxygens in MD simulations are enclosed in colored boxes. Occupancy of water oxygen is displayed by isosurface. Gray regions represent 50% occupancy and the red regions represent 75% occupancy.

By building volumetric maps, the average positions of waters during MD simulations could be compared with the water positions in crystal structures. Waters 400, 401 and 402 are part of the H-bond network connecting SB216, D85 and D212 within CC; while waters 400, 401, 403, 406, 407, 408 and 409 tend to participate in proton wires between the CC and the EC. In the CC, the upward orientation of R82 side-chain in t-bR, c-bR, t-M1 and c-M1 models causes water 400 to 403 to be more stably occupied than in late stage where the density of these water shifts downward with R82. Notably, water 401 is destabilized (below 50% occupancy) in c-O and t-O due to competition from the strong H-bonds formed between SB216 and D212 (Table 2). Only when D212 is protonated in t-O’, is occupancy of water in position 401 restored. In the EC, waters in position 406 to 409 became more stable when the R82 side-chain orients downward in c-M2, c-O, t-O and t-O’ models. Therefore, the side-chain motion of R82 induces the repositioning of waters between the CC and the EC, which further modifies the proton transfer pathway. Waters rearrange in ways that can be functionally significant. In t-O there is a high probability of forming direct H- bonds from SB216 to D212 and from D85 to D212. By contrast, in the rest of the models, D85 and D212 are separated by at least one water molecule (Figure 4 and Table S1). Also in t-O, water in position 401 shifts toward to position 400 where it can participate in a proton wire with waters in position 403, 407 and 409. The water in these positions tend to facilitate the proton transfer along D85-D212-W400-W403-(R82)-W407-(W409)-E204 that could convert t-O to t-bR state. Nevertheless, the existence of a H-bonded chain is necessary but not sufficient for proton transfer, which requires the system to surpass the energy barrier for moving a proton from the CC to the EC. Given the flexible motions of the R82 side-chain and waters, proton transfer pathways between the CC and the EC are more complex than proposed for the cytoplasmic half channel.^22^

**Possible role of R82 in proton transfer between the CC and the EC.** Although R82 appears to be important in proton pumping, several R82 mutants can still pump protons, though with decreased efficiency.^69,70^ Mutated protein substituting R82 with Gln,^69,70^ Ala^71^ or Lys^13^ delay the proton release from the EC until after proton uptake via D96.^72^ In the R82H mutant,^73^ the t-O to t-bR transition is slowed, indicating that R82 is important for the proton transfer from the CC to the EC. The analysis here will focus on the wild-type protein where the large, positively charged R82 appears to be a major component in the region between the CC and the EC.

Throughout the photocycle, the R82 side-chain changes its orientation pointing toward the CC or the EC as seen in crystal structures trapped in different intermediates^66^ and in the MD trajectories presented here (Figure 2). The motion of R82 is associated with reorganization of H-bond and salt bridge interactions with waters and acidic residue side-chains. It has been proposed^33,57,74^ that R82 would first donate a proton to the EC and then move upward in its neutral form to be reprotonated by D85 via a proton hopping mechanism.

Proton transfer in proteins is often assumed to adopt a Grotthus mechanism, which relies on groups such as water, hydroxyls and protonated carboxylic acids. The possession of lone pairs of electrons allows these moieties to function simultaneously as a proton donor and acceptor. By contrast, the protonated arginine side-chain cannot accept a proton due to the conjugation between the double bond and the nitrogen lone pair.

To study the possible role of an Arg side chain in facilitating proton transfer, quantum mechanical IRC calculations were performed on a simplified model containing only a methyl guanidinium, hydronium and hydroxide ions. This provides a strong driving force to transfer a proton from the hydronium to the hydroxide through the guanidinium (Figure 5) without considering the electrostatic interactions or other H-bonds in the protein. In all cases of guanidinium-facilitated proton transfer, the hydroxide first accepts a proton from a guanidinium N, which then accepts a proton from the adjacent hydronium. It was found that if the hydroxide and hydronium oxygens were initially within H-bond distance of the same amine, a proton hopped from a guanidinium N to the hydroxide, and then another proton hopped from the hydronium to the guanidinium N, resulting in methyl-guanidinium and two waters (Figure 5 A to D, Movie S1A). Conversely, if the hydroxide and hydronium oxygens were initially within H-bond distance of different amines, then proton donation and acceptance took place at different N’s, forming a high-energy tautomer (Figure 5E to H) that are unable to convert back to the low energy guanidinium tautomer. One exception is when hydroxide and hydronium ions were placed on the same side of Ns-Cζ axis, near Nη2 and Ns respectively (Figure 5D and Movie S1B). After the hydroxide ion accepts a proton from Ns, the newly generated water forms a Zundel ion with the hydronium, and then returns the proton to the Nε leaving Nη2 intact.

**Figure 5.**
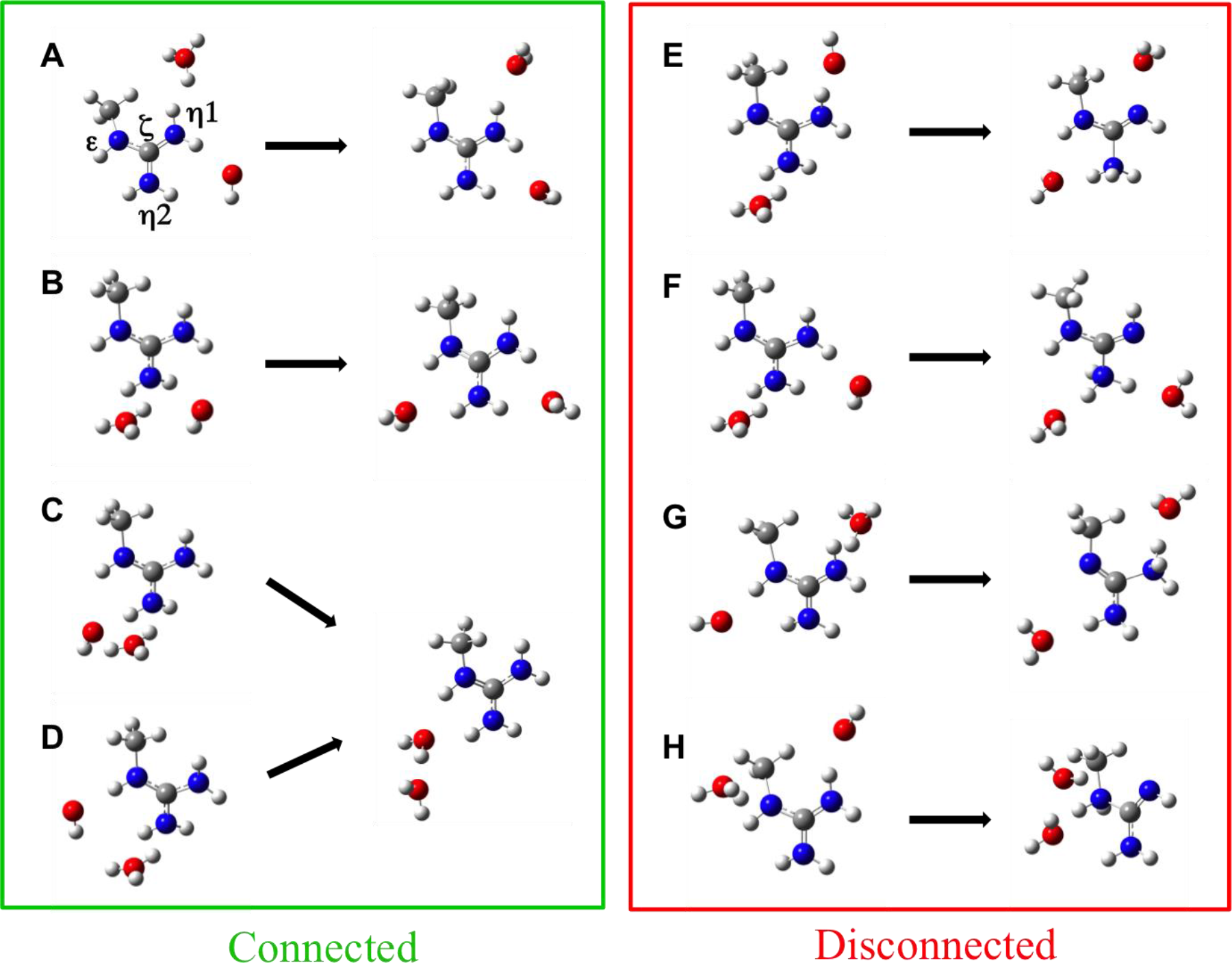
IRC reaction paths starting with hydroxide and hydronium ions around methyl guanidinium in different initial positions. The left panel shows four different initial geometries that can transfer a proton through guanidinium successfully; the right panel shows four cases that lead to high-energy guanidinium tautomers.

The IRC calculations indicate that R82 can participate in proton transfer via proton hopping if the initial geometry allows proton donation and acceptance by the same amine. The first step of this process is to generate a deprotonated R82 intermediate. A recent NMR study^75^ revisited the pKa of the arginine side-chain in water, placing it higher than 13. However, a transiently deprotonated arginine side-chain may still play a role as an unstable intermediate in the proton transfer from D85 to the EC, which takes place on the millisecond timescale. With a pKa value about 12 to 13, the free energy barrier for neutralizing an arginine side-chain in solution is ΔG = -2.303*RT*pKa, which equals -16.4 to -17.7kcal/mol. IR studies suggest that the proton affinity of R82 may be lower in situ than that of an arginine in solution, which would lower the barrier.^76^

Table 3 describes proton wires formed within a single MD snapshot that connect a given side-chain N of R82 to a residue of the CC or the EC. As R82 remains protonated in the MD trajectories, the reorientation of these paths to accommodate a transiently deprotonated Arg were not considered here. The side-chain of R82 is likely to be connected to D212 via Nrj1 (>90% in t-bR, c-bR, t-M1 and c-M1 states, decreasing from 70% to 12% in c-M2, c-O, t-O and t-O’ states), or via Nη2 less frequently (30∽67% in t-bR, c-bR, t-M1 and c-M1). The connection from R82 to D85 is only possible via Nrj1, which becomes less frequent along the reaction cycle. The connections from R82 to the EC are found via Nη2 in all the states, or via Nη1 in c-M2, c-O, t-O and t-O’ states (>80%). The R82-originated proton wires show two typical patterns representing the early stage (t-bR, c-bR, t-M1 and c-M1) and the late stage (cM2, c-O, t-O and t-O’) respectively (Figure 3B).

Changes in propensity of different proton wires originating from the R82 side-chain suggest that the stability of the connections varies in different intermediates. The IRC study implies that the R82 side-chain must be deprotonated first to create a transient “proton hole” for proton transfer by hopping. A proton-wire with longer lifetime in the trajectories is more likely to have an elevated turnover number for creating a transient R82. The direction of proton transfer could be inferred by comparing the probability of forming proton wires from the same R82 side-chain N to the CC and the EC side with less than six sequential H-bonds (Table S2). For instance, in c-M2, c-O, t-O and t-O’ models, each R82 side-chain N atom forms proton-wires to an EC residue with a high probability (>80%); while proton wires connect R82 to the CC side only via Nη1 with decreasing probability (from 70% to 12%) along the reaction cycle. Thus, proton transfer favors the CC to EC direction in these states. By contrast, in t-bR, c-bR, c-M1 and t-M1 models, the probabilities of forming these proton wires on the two side are indistinguishable, *i.e.,* R82 connects to the CC via Nη 1, to the EC via Nε, and to both the CC and the EC with roughly equal probabilities. The proton transfer between the CC and the EC in the early states (t-bR, c-bR, c-M1 and t-M1) tends to be energetically unfavorable due to the relative proton affinities of the two clusters.^33^ Therefore, the relative propensity of proton wires in the two direction via R82 suggest a higher probability of proton transfer from the CC to the EC in t-O and t-O’ as required for the reaction cycle (Figure 3).

**Alternative pathways for proton translocation from the CC to the EC.** QM calculations indicate that proton transfer can propagate though water wires in a quasi-concerted manner over a few water molecules.^77–80^ Time-resolved FTIR and in situ H2^18^O/H2^16^O exchange FTIR experiments suggest that the controlled Grotthuss proton transfer is more likely to take place in bacteriorhdopsin than random proton migration that occurs in liquid water.^81^ With the positions of the high-occupancy waters found in the MD trajectories, we searched for proton wires offering pathways for proton translocation between the CC and the EC that do not involve R82. Multistep proton wires connecting a protonated CC Aspartate to the EC Glutamates in a single snapshot were found in 0.02% and 0.3% of the MD frames in c-M2 and c-O models respectively (Table 3). Although rare, such a proton-conducting pathway once available may permit more efficient proton transfer with a lower barrier than proton hopping via a deprotonated R82. Since a particularly stable H-bond (80% probability) was found between D85 and D212 in the t-O state, a t-O’ state with D85 proton transferred to D212 was explored. The existence of this transient t-O’ state is supported by FTIR experiments^82^ and QM/MM simulations.^83^ Transient D212 protonation is also found in the c-M1 state by continuum electrostatic calculation.^39^ Remarkably, the t-O’ state model increased the propensity of forming proton wires from D212 to E204 by nine fold (2.7%), indicating that a transient t-O’ state could enhance the Grotthuss-type proton transfer from the CC to the EC during the t-O to t-bR transition.

The water-facilitated proton wires are only oriented to transfer protons from the CC to the EC, but do not form in the reverse direction, even when the EC residues could act as donors in the bR or M1 states. These results also imply that proton transfer pathway from the CC to the EC is accessible either before (c-M2) or more likely after (c-O and t-O’) the proton uptake from the N-side to the CC. In c-M2 state, there is only one proton in the CC (on D85) which gives rise to a -1 total charge of the CC, while the downward oriented R82 partially neutralizes the double-deprotonated EC, hence, proton transfer from the CC to the EC in c-M2 lacks an electrostatic driving force. The observed proton wires were usually composed of four to six waters or polar side-chains between the CC and the EC, forming five to seven sequential H-bonds (here we call them steps), with the majority having six steps (Table S1). Several representative snapshots shown in Figure 6 indicate that besides water, Y57 and T205 side-chains also participate in some of the proton-transfer pathways.

**Figure 6.**
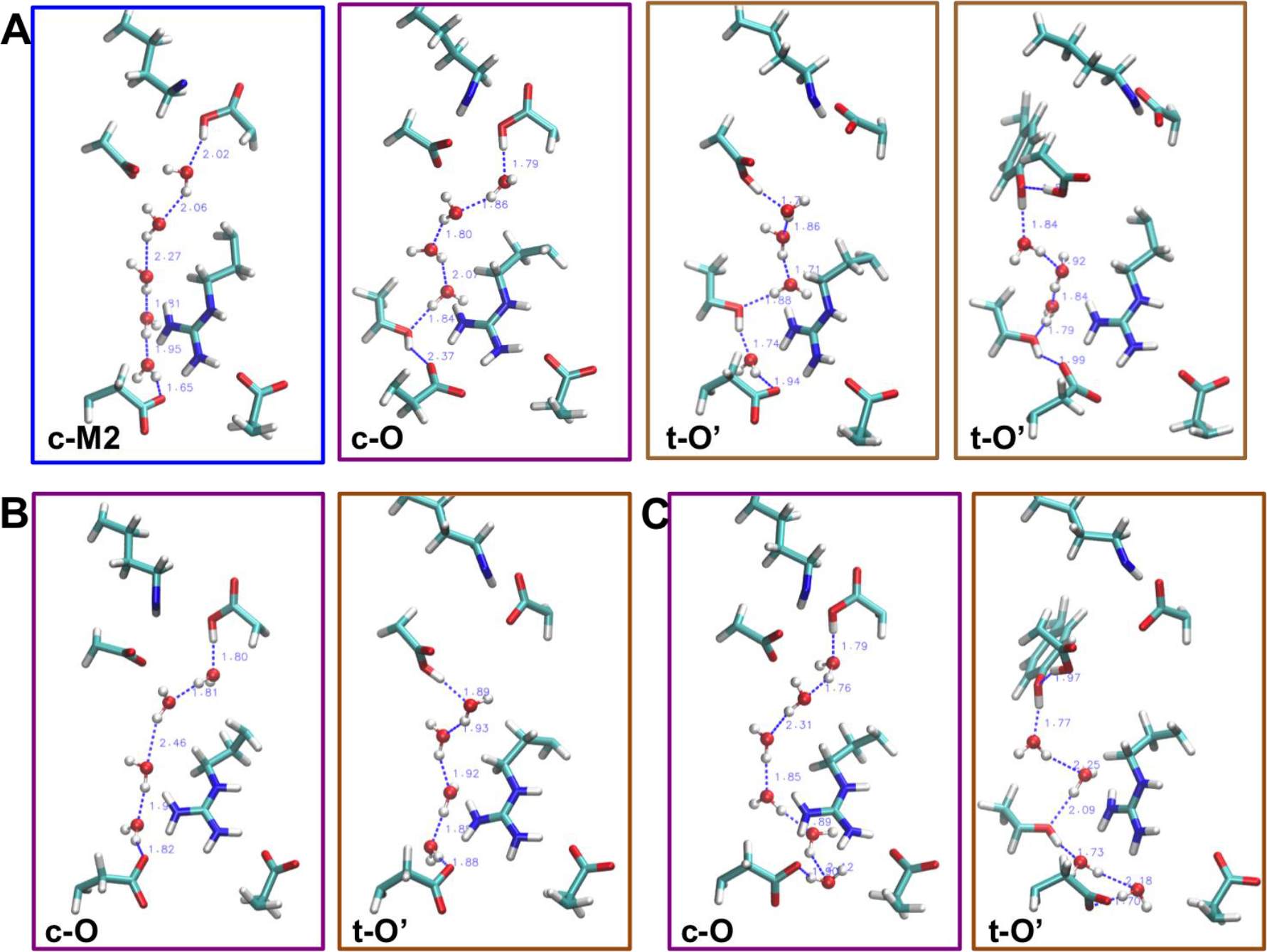
Examples of complete proton wires connecting the CC to the EC. Panel A is for paths requiring six steps. Panel B and C show paths made by five or seven steps respectively.

The Grotthuss mechanism has been revisited in recent *ab initio* MD simulations of water wires,^79^ which suggest that proton transfer is a process involving a broader distribution of pathways and timescales. Therefore, although the Grotthuss-type proton transfer was thought to occur in picosecond timescale,^48,49^ forming such a pre-transfer geometry requires pre-solvation and prearrangement of all groups on the pathway, which could take place in much longer timescale. Depending on the length and nature of the pathway, this process may occur in nanoseconds to milliseconds. Most of the proton transfer steps in bacteriorhodopsin photocycle are slow processes, occurring in microseconds to milliseconds.^2,4,32^ Proton translocation in the extracellular half-channel is the slowest step, typically occurring in a few milliseconds at neutral pH. This process is likely to be slowed by generating a high-energy neutral Arg if following a proton-hopping pathway, or because a lower-energy but longer Grotthuss-type pathway is rarely formed. Since R82 does not actively participate in a Grotthuss-type pathway, this may help explain how the R82 mutant proteins can still pump protons.^13,69–71,73^

## Conclusion

Computational models of bacteriorhodopsin in four intermediate states t-bR, c-M1, c-M2 and t-O as well as four hypothetical states c-bR, t-M1, c-O and t-O’ were constructed to explore the proton transfer pathways in the extracellular half-channel. The equilibrium distribution of R82, a residue that lies between the CC and the EC, was found correlated with the protonation states of the amino acids in the EC. R82 oscillates between the CC and the EC during the early stage of the proton-pumping cycle when the EC has a net charge of**-**1 in t-bR, c-bR, t-M1 and c-M1 states, but adopts a downward orientation when the EC charge is **-**2 in c-M2, c-O, t-O and t-O’ states. The residue motions of the two EC glutamates are also highly correlated with their protonation states and the R82 orientation. Volumetric maps of water occupancy in equilibrated trajectories suggest distinct water-residue interactions in different states of the protein. These findings are consistent with previous crystallographic and simulation studies of a few intermediated states. Furthermore, the correlated motions of R82 and waters reflect the alternation of H-bond network and consequently the possible pathways for proton transfer in each step of the proton-pumping cycle.

Quantum mechanical IRC calculations show the geometrical restrictions for the proton transfer reaction involving an arginine, *i.e.,* an initial geometry allowing proton donation and acceptance by the same amine. A proton-hopping mechanism is possibly adopted where the proton is first transferred from R82 to the acceptor forming a transiently deprotonated R82, which then accepts a proton from the proton donor. In all the models, R82 can form proton wires to both the CC and the EC via the same side-chain N. Therefore, The CC and the EC can be connected via R82 in all stages of the reaction cycle, which leaves the protein susceptible to “wrong-way” proton transfer. Thus, other barriers to proton transfer and the relative proton affinity of the proton donors and acceptors must control the proton translocation in this region. The probabilities of forming these proton wires from R82 to either the CC or the EC are indistinguishable in t-bR, c-bR, c-M1 and t-M1 models. By contrast, in c-M2, c-O, t-O and t-O’ states, the proton wires connecting R82 to the EC dominate; the lifetime of proton wires connecting R82Nη1 to the EC increase while that to the EC decreases. This trend does lead to a preferred proton transfer direction from the CC to the EC in the later states as required for pumping. Although, there are hydrogen bonded connections between the CC and the EC via R82 in all states, proton transfer via this route requires forming a high-energy neutral arginine, which will elevate the reaction barrier and thus decrease the proton transfer rate.

Based on the H-bond and proton wire analysis, alternative Grotthuss-type pathways were found between the CC and the EC, solely through waters and side-chains of polar residues excluding R82. This could help explain why R82 mutant proteins retain the proton-pumping ability. In the extracellular half-channel, the Grotthuss-type transfer only occurs vectorially in the late stage, with an enhanced probability in a t-O’ transient state where the proton on D85 has been transferred to D212. These findings support active proton translocation from the CC to the EC in the appropriate stage of the reaction cycle during the transition of t-O to t-bR ground state.

Taking together, this MD simulation-based study revealed novel aspects of long-range proton-translocation within the extracellular half-channel of bacteriorhodopsin. Nevertheless, there is the possibility that such kind of brute force simulations did not fully converge. To achieve sufficient sampling and ensure convergence, we envisage a future direction of applying enhanced sampling methods to the bacteriorhodopsin models of interest.

## Acknowledgements

This research was supported by the National Science Foundation (NSF) Grant number MCB1519640. The computations were performed on Texas Advanced Computing Center (TACC) cluster Stampede and storage Ranch. The TACC resources were provided through the Startup Allocation MCB140055 of the Extreme Science and Engineering Discovery Environment (XSEDE), which is supported by NSF grant number ACI-1053575. This research was supported, in part, by a grant of computer time from the City University of New York High Performance Computing Center under NSF Grants CNS-0855217 and CNS-0958379, as well as National Institute on Minority Health and Health Disparities Grant 8G12MD7603-28 for infrastructure. The authors are grateful to Dr. Qiang Cui for providing the initial coordinates of one simulation model as a reference, and to Dr. Benoit Roux and Dr. Yen-lin Lin for counseling on compute resources.

## References

1. Gunner MR, Amin M, Zhu X, Lu J. Molecular mechanisms for generating transmembrane proton gradients. Biochimica et biophysica acta 2013; 1827 (8-9): 892–913.

2. Balashov SP. Protonation reactions and their coupling in bacteriorhodopsin. Biochim Biophys Acta 2000; 1460(1):75–94.

3. Wraight CA. Chance and design--proton transfer in water, channels and bioenergetic proteins. Biochimica et biophysica acta 2006; 1757(8): 886–912.

4. Lanyi JK. Proton transfers in the bacteriorhodopsin photocycle. Biochimica et biophysica acta 2006; 1757(8): 1012–1018.

5. Braiman MS, Bousche O, Rothschild KJ. Protein dynamics in the bacteriorhodopsin photocycle: submillisecond Fourier transform infrared spectra of the L, M, and N photointermediates. Proc Natl Acad Sci USA 1991; 88(6): 2388–2392.

6. Braiman MS, Mogi T, Marti T, Stern LJ, Khorana HG, Rothschild KJ. Vibrational spectroscopy of bacteriorhodopsin mutants: light-driven proton transport involves protonation changes of aspartic acid residues 85, 96, and 212. Biochemistry 1988; 27(23): 8516–8520.

7. Phatak P, Ghosh N, Yu HB, Cui Q, Elstner M. Amino acids with an intermolecular proton bond as proton storage site in bacteriorhodopsin. P Natl Acad Sci USA 2008; 105(50): 19672–19677.

8. Hessling B, Herbst J, Rammelsberg R, Gerwert K. Fourier transform infrared double-flash experiments resolve bacteriorhodopsin’s M1 to M2 transition. Biophys J 1997; 73(4): 2071–2080.

9. Gerwert K, Hess B, Soppa J, Oesterhelt D. Role of aspartate-96 in proton translocation by bacteriorhodopsin. Proc Natl Acad Sci USA 1989; 86: 4943–4947.

10. Callender R, Deng H. Nonresonance Raman difference spectroscopy: a general probe of protein structure, ligand binding, enzymatic catalysis, and the structures of other biomacromolecules. Annu Rev Biophys Biomol Struct 1994; 23: 215–245.

11. Hatanaka M, Sasaki J, Kandori H, Ebrey TG, Needleman R, Lanyi JK, Maeda A. Effects of arginine-82 on the interactions of internal water molecules in bacteriorhodopsin. Biochemistry 1996; 35(20): 6308–6312.

12. Maeda A, Kandori H, Yamazaki Y, Nishimura S, Hatanaka M, Chon YS, Sasaki J, Needleman R, Lanyi JK. Intramembrane signaling mediated by hydrogen-bonding of water and carboxyl groups in bacteriorhodopsin and rhodopsin. J Biochem 1997; 121(3): 399–406.

13. Balashov SP, Govindjee R, Imasheva ES, Misra S, Ebrey TG, Feng Y, Crouch RK, Menick DR. The 2 Pk(a) of Aspartate-85 and Control of Thermal-Isomerization and Proton Release in the Arginine-82 to Lysine Mutant of Bacteriorhodopsin. Biochemistry 1995; 34(27): 8820–8834.

14. Lanyi JK. Progress toward an explicit mechanistic model for the light-driven pump, bacteriorhodopsin. FEBS letters 1999; 464(3): 103–107.

15. Kandt C, Schlitter J, Gerwert K. Dynamics of water molecules in the bacteriorhodopsin trimer in explicit lipid/water environment. Biophys J 2004; 86(2): 705–717.

16. Belrhali H, Nollert P, Royant A, Menzel C, Rosenbusch JP, Landau EM, Pebay-Peyroula E. Protein, lipid and water organization in bacteriorhodopsin crystals: a molecular view of the purple membrane at 1.9 A resolution. Structure 1999; 7(8): 909–917.

17. Schobert B, Brown LS, Lanyi JK. Crystallographic structures of the M and N intermediates of bacteriorhodopsin: assembly of a hydrogen-bonded chain of water molecules between Asp-96 and the retinal Schiff base. J Mol Biol 2003; 330(3): 553–570.

18. Sass HJ, Buldt G, Gessenich R, Hehn D, Neff D, Schlesinger R, Berendzen J, Ormos P. Structural alterations for proton translocation in the M state of wild-type bacteriorhodopsin. Nature 2000; 406 (6796): 649–653.

19. Okumura H, Murakami M, Kouyama T. Crystal structures of acid blue and alkaline purple forms of bacteriorhodopsin. J Mol Biol 2005; 351(3): 481–495.

20. Zhang J, Yamazaki Y, Hikake M, Murakami M, Ihara K, Kouyama T. Crystal structure of the O intermediate of the Leu93-->Ala mutant of bacteriorhodopsin. Proteins 2012; 80(10): 2384–2396.

21. Wickstrand C, Dods R, Royant A, Neutze R. Bacteriorhodopsin: Would the real structural intermediates please stand up? Biochimica et biophysica acta 2015; 1850(3): 536–553.

22. Freier E, Wolf S, Gerwert K. Proton transfer via a transient linear water-molecule chain in a membrane protein. P Natl Acad Sci USA 2011; 108(28): 11435–11439.

23. Wang T, Sessions AO, Lunde CS, Rouhani S, Glaeser RM, Duan Y, Facciotti MT. Deprotonation of D96 in Bacteriorhodopsin Opens the Proton Uptake Pathway. Structure 2013; 21(2): 290–297.

24. Warshel A, Chu ZT. Nature of the surface crossing process in bacteriorhodopsin: Computer simulations of the quantum dynamics of the primary photochemical event. J Phys Chem B 2001; 105(40): 9857–9871.

25. Braun-Sand S, Warshel A. Simulating the primary proton transport event in bacteriorhodopsin. Biophysical Journal 2005; 88(1): 507a–507a.

26. Bondar AN, Suhai S, Fischer S, Smith JC, Elstner M. Suppression of the back proton-transfer from Asp85 to the retinal Schiff base in bacteriorhodopsin: A theoretical analysis of structural elements. Journal of structural biology 2007; 157(3): 454–469.

27. Bondar AN, Elstner M, Suhai S, Smith JC, Fischer S. Mechanism of primary proton transfer in bacteriorhodopsin. Structure 2004; 12(7): 1281–1288.

28. Braun-Sand S, Sharma PK, Chu ZT, Pisliakov AV, Warshel A. The energetics of the primary proton transfer in bacteriorhodopsin revisited: It is a sequential light-induced charge separation after all. Bba-Bioenergetics 2008; 1777(5): 441–452.

29. Goyal P, Ghosh N, Phatak P, Clemens M, Gaus M, Elstner M, Cui Q. Proton Storage Site in Bacteriorhodopsin: New Insights from Quantum Mechanics/Molecular Mechanics Simulations of Microscopic pK(a) and Infrared Spectra. Journal of the American Chemical Society 2011; 133(38): 14981–14997.

30. Wolf S, Freier E, Gerwert K. A Delocalized Proton-Binding Site within a Membrane Protein. Biophysical Journal 2014; 107(1): 174–184.

31. Wolf S, Freier E, Gerwert K. How Does a Membrane Protein Achieve a Vectorial Proton Transfer Via Water Molecules? Chemphyschem 2008; 9(18): 2772–2778.

32. Lorenz-Fonfria VA, Kandori H. Spectroscopic and kinetic evidence on how bacteriorhodopsin accomplishes vectorial proton transport under functional conditions. Journal of the American Chemical Society 2009; 131(16): 5891–5901.

33. Onufriev A, Smondyrev A, Bashford D. Affinity changes driving unidirectional proton transport in the bacteriorhodopsin photocycle. J Mol Biol 2003; 332(5): 1183–1193.

34. Dencher NA, Heyn MP. Bacteriorhodopsin monomers pump protons. FEBS letters 1979; 108(2): 307–310.

35. The PyMOL Molecular Graphics System, Version 1.5.0.4 Schrödinger, LLC.

36. Phillips JC, Braun R, Wang W, Gumbart J, Tajkhorshid E, Villa E, Chipot C, Skeel RD, Kale L, Schulten K. Scalable molecular dynamics with NAMD. J Comput Chem 2005; 26(16): 1781–1802.

37. MacKerell AD, Bashford D, Bellott M, Dunbrack RL, Evanseck JD, Field MJ, Fischer S, Gao J, Guo H, Ha S, Joseph-McCarthy D, Kuchnir L, Kuczera K, Lau FTK, Mattos C, Michnick S, Ngo T, Nguyen DT, Prodhom B, Reiher WE, Roux B, Schlenkrich M, Smith JC, Stote R, Straub J, Watanabe M, Wiorkiewicz-Kuczera J, Yin D, Karplus M. All-atom empirical potential for molecular modeling and dynamics studies of proteins. J Phys Chem B 1998; 102(18): 3586–3616.

38. Jorgensen WL, Jenson C. Temperature dependence of TIP3P, SPC, and TIP4P water from NPT Monte Carlo simulations: Seeking temperatures of maximum density. J Comput Chem 1998; 19(10): 1179–1186.

39. Song Y, Mao J, Gunner MR. Calculation of proton transfers in bacteriorhodopsin bR and M intermediates. Biochemistry 2003; 42(33): 9875–9888.

40. Ryckaert JP, Ciccotti G, Berendsen HJC. Numerical-Integration of Cartesian Equations of Motion of a System with Constraints - Molecular-Dynamics of N-Alkanes. J Comput Phys 1977; 23(3): 327–341.

41. Humphrey W, Dalke A, Schulten K. VMD: Visual molecular dynamics. Journal of molecular graphics & modelling 1996; 14(1): 33–38.

42. Maupin CM, Saunders MG, Thorpe IF, McKenna R, Silverman DN, Voth GA. Origins of enhanced proton transport in the Y7F mutant of human carbonic anhydrase II. Journal of the American Chemical Society 2008; 130(34): 11399–11408.

43. Arnarez C, Mazat JP, Elezgaray J, Marrink SJ, Periole X. Evidence for cardiolipin binding sites on the membrane-exposed surface of the cytochrome bc1. Journal of the American Chemical Society 2013; 135(8): 3112–3120.

44. Raunest M, Kandt C. Locked on one side only: ground state dynamics of the outer membrane efflux duct TolC. Biochemistry 2012; 51(8): 1719–1729.

45. Bondi A. van der Waals Volumes and Radii. J Phys Chem-Us 1964; 68(3): 441–451.

46. Alvarez S. A cartography of the van der Waals territories. Dalton T 2013; 42(24): 8617–8636.

47. Jeffrey GA. An Introduction to Hydrogen Bonding. New York: Oxford University Press.; 1997.

48. Agmon N. The Grotthuss Mechanism. Chem Phys Lett 1995; 244 (5–6): 456–462.

49. Cukierman S. Et tu, Grotthuss! and other unfinished stories. Bba-Bioenergetics 2006; 1757(8): 876–885.

50. Fukui K, Kato S, Fujimoto H. Constituent Analysis of Potential Gradient Along a Reaction Coordinate - Method and an Application to Ch4 + T Reaction. Journal of the American Chemical Society 1975; 97(1): 1–7.

51. Ishida K, Morokuma K, Komornicki A. Intrinsic Reaction Coordinate - an Abinitio Calculation for Hnc-]Hcn and H-+Ch4-]Ch4+H-. J Chem Phys 1977; 66(5): 2153–2156.

52. Frisch MJT, G. W.; Schlegel, H. B.; Scuseria, G. E.; Robb, M. A.; Cheeseman, J. R.; Scalmani, G.; Barone, V.; Mennucci, B.; Petersson, G. A.; Nakatsuji, H.; Caricato, M.; Li, X.; Hratchian, H. P.; Izmaylov, A. F.; Bloino, J.; Zheng, G.; Sonnenberg, J. L.; Hada, M.; Ehara, M.; Toyota, K.; Fukuda, R.; Hasegawa, J.; Ishida, M.; Nakajima, T.; Honda, Y.; Kitao, O.; Nakai, H.; Vreven, T.; Montgomery, J. A., Jr.; Peralta, J. E.; Ogliaro, F.; Bearpark, M.; Heyd, J. J.; Brothers, E.; Kudin, K. N.; Staroverov, V. N.; Kobayashi, R.; Normand, J.; Raghavachari, K.; Rendell, A.; Burant, J. C.; Iyengar, S. S.; Tomasi, J.; Cossi, M.; Rega, N.; Millam, M. J.; Klene, M.; Knox, J. E.; Cross, J. B.; Bakken, V.; Adamo, C.; Jaramillo, J.; Gomperts, R.; Stratmann, R. E.; Yazyev, O.; Austin, A. J.; Cammi, R.; Pomelli, C.; Ochterski, J. W.; Martin, R. L.; Morokuma, K.; Zakrzewski, V. G.; Voth, G. A.; Salvador, P.; Dannenberg, J. J.; Dapprich, S.; Daniels, A. D.; Farkas, Ö.; Foresman, J. B.; Ortiz, J. V.; Cioslowski, J.; Fox, D. J. Gaussian 09, Revision D.01. Wallingford CT: Gaussian, Inc.; 2009.

53. Zhang JH, Wang YC, Jiao HJ, Geng ZY. On the gas-phase OsOn+ (n=0–3) catalyzed reduction of N2O by H-2: A density functional study. Comput Theor Chem 2014; 1042: 35–40.

54. Jeon IS, Ahn DS, Park SW, Lee S, Kim B. Structures and isomerization of neutral and zwitterion serine-water clusters: Computational study. Int J Quantum Chem 2005; 101(1): 55–66.

55. Inostroza-Rivera R, Herrera B, Toro-Labbe A. Using the reaction force and the reaction electronic flux on the proton transfer of formamide derived systems. Phys Chem Chem Phys 2014; 16(28): 14489–14495.

56. Spassov VZ, Luecke H, Gerwert K, Bashford D. pKa calculations suggest storage of an excess proton in a hydrogen- bonded water network in bacteriorhodopsin. J Mol Biol 2001; 312(1): 203–219.

57. Gunner MR, Mao J, Song Y, Kim J. Factors influencing energetics of electron and proton transfers in proteins. What can be learned from calculations. Biochim Biophys Acta 2006; 1757: 942–968.

58. Song Y, Gunner MR. Halorhodopsin pumps Cl- and bacteriorhodopsin pumps protons by a common mechanism that uses conserved electrostatic interactions. Proc Natl Acad Sci USA 2014; 111(46): 16377–16382.

59. Bajaj VS, Mak-Jurkauskas ML, Belenky M, Herzfeld J, Griffin RG. Functional and shunt states of bacteriorhodopsin resolved by 250 GHz dynamic nuclear polarization-enhanced solid-state NMR. Proc Natl Acad Sci USA 2009; 106(23): 9244–9249.

60. Nina M, Roux B, Smith JC. Functional Interactions in Bacteriorhodopsin - a Theoretical-Analysis of Retinal Hydrogen-Bonding with Water. Biophysical Journal 1995; 68(1): 25–39.

61. Tajkhorshid E, Baudry J, Schulten K, Suhai S. Molecular dynamics study of the nature and origin of retinal’s twisted structure in bacteriorhodopsin. Biophysical Journal 2000; 78(2): 683–693.

62. Babitzki G, Denschlag R, Tavan P. Polarization effects stabilize bacteriorhodopsin’s chromophore binding pocket: a molecular dynamics study. J Phys Chem B 2009; 113(30): 10483–10495.

63. Bondar AN, Knapp-Mohammady M, Suhai S, Fischer S, Smith JC. Ground-state properties of the retinal molecule: from quantum mechanical to classical mechanical computations of retinal proteins. Theor Chem Acc 2011; 130 (4–6): 1169–1183.

64. Vanommeslaeghe K, Hatcher E, Acharya C, Kundu S, Zhong S, Shim J, Darian E, Guvench O, Lopes P, Vorobyov I, MacKerell AD. CHARMM General Force Field: A Force Field for Drug-Like Molecules Compatible with the CHARMM All-Atom Additive Biological Force Fields. J Comput Chem 2010; 31(4): 671–690.

65. Clemens M, Phatak P, Cui Q, Bondar AN, Elstner M. Role of Arg82 in the Early Steps of the Bacteriorhodopsin Proton-Pumping Cycle. J Phys Chem B 2011; 115(21): 7129–7135.

66. Lanyi JK, Schobert B. Mechanism of proton transport in bacteriorhodopsin from crystallographic structures of the K, L, M-1, M-2, and M-2 ‘ intermediates of the photocycle. J Mol Biol 2003; 328(2): 439–450.

67. Leontyev I, Stuchebrukhov A. Accounting for electronic polarization in non-polarizable force fields. Phys Chem Chem Phys 2011; 13(7): 2613–2626.

68. Bondar AN, Baudry J, Suhai S, Fischer S, Smith JC. Key role of active-site water molecules in bacteriorhodopsin proton-transfer reactions. J Phys Chem B 2008; 112(47): 14729–14741.

69. Otto H, Marti T, Holz M, Mogi T, Stern LJ, Engel F, Khorana HG, Heyn MP. Substitution of Amino-Acids Asp-85, Asp-212, and Arg-82 in Bacteriorhodopsin Affects the Proton Release Phase of the Pump and the Pk of the Schiff-Base. P Natl Acad Sci USA 1990; 87(3): 1018–1022.

70. Govindjee R, Misra S, Balashov SP, Ebrey TG, Crouch RK, Menick DR. Arginine-82 regulates the pK(a) of the group responsible for the light-driven proton release in bacteriorhodopsin. Biophysical Journal 1996; 71(2): 1011–1023.

71. Balashov SP, Govindjee R, Kono M, Imasheva E, Lukashev E, Ebrey TG, Crouch RK, Menick DR, Feng Y. Effect of the Arginine-82 to Alanine Mutation in Bacteriorhodopsin on Dark-Adaptation, Proton Release, and the Photochemical Cycle. Biochemistry 1993; 32(39): 10331–10343.

72. Balashov SP, Lu M, Imasheva ES, Govindjee R, Ebrey TG, Othersen B, Chen YM, Crouch RK, Menick DR. The proton release group of bacteriorhodopsin controls the rate of the final step of its photocycle at low pH. Biochemistry 1999; 38(7): 2026–2039.

73. Imasheva ES, Balashov SP, Ebrey TG, Chen N, Crouch RK, Menick DR. Two groups control light-induced Schiff base deprotonation and the proton affinity of Asp(85) in the Arg(82) his mutant of bacteriorhodopsin. Biophysical Journal 1999; 77(5): 2750–2763.

74. Brown L, Bonet L, Needleman R, Lanyi J. Estimated acid dissociation constants of the Schiff base, Asp-85, and Arg-82 during the bacteriorhodopsin photocycle. Biophys J 1993; 65(1): 124–3226.

75. Fitch CA, Platzer G, Okon M, Garcia-Moreno EB, McIntosh LP. Arginine: Its pKa value revisited. Protein Sci 2015; 24(5): 752–761.

76. Hutson MS, Alexiev U, Shilov SV, Wise KJ, Braiman MS. Evidence for a perturbation of arginine-82 in the bacteriorhodopsin photocycle from time-resolved infrared spectra. Biochemistry 2000; 39(43): 13189–13200.

77. Pomes R, Roux B. Free energy profiles for H+ conduction along hydrogen-bonded chains of water molecules. Biophysical Journal 1998; 75(1): 33–40.

78. Cui Q, Karplus M. Is a “proton wire” concerted or stepwise? A model study of proton transfer in carbonic anhydrase. J Phys Chem B 2003; 107(4): 1071–1078.

79. Hassanali A, Giberti F, Cuny J, Kuhne TD, Parrinello M. Proton transfer through the water gossamer. Proc Natl Acad Sci USA 2013; 110(34): 13723–13728.

80. Kaila VRI, Hummer G. Energetics and dynamics of proton transfer reactions along short water wires. Phys Chem Chem Phys 2011; 13(29): 13207–13215.

81. Garczarek F, Gerwert K. Functional waters in intraprotein proton transfer monitored by FTIR difference spectroscopy. Nature 2006; 439 (7072): 109–112.

82. Dioumaev AK, Brown LS, Needleman R, Lanyi JK. Fourier transform infrared spectra of a late intermediate of the bacteriorhodopsin photocycle suggest transient protonation of Asp-212. Biochemistry 1999; 38(31): 10070–10078.

83. Phatak P, Frahmcke JS, Wanko M, Hoffmann M, Strodel P, Smith JC, Suhai S, Bondar AN, Elstner M. Long-Distance Proton Transfer with a Break in the Bacteriorhodopsin Active Site. Journal of the American Chemical Society 2009; 131(20): 7064–7078.

